# Ultrastructural changes at auditory nerve synapses following moderate noise exposure

**DOI:** 10.64898/2025.12.30.697061

**Authors:** Denesha Gorman, Nicole F. Wong, Colin W. Schupbach, Stacy L. DiCenso, Sophie C. Xu-Friedman, Kevin M. Boergens, Amanda M. Lauer, Angeles Salles, Matthew A. Xu-Friedman

## Abstract

Moderate noise exposure is a common experience, yet its impact on central auditory synapses remains poorly understood. We study this issue at the first synapses in the central auditory pathway formed by auditory nerve afferents onto bushy cells in the cochlear nucleus, called endbulbs of Held. Non-traumatic noise exposure alters endbulb properties, decreasing the probability of vesicle release and enlarging the pool of releasable vesicles as assessed using electrophysiological methods and immunolabelling. These changes appear homeostatic, to maintain synaptic efficacy during periods of high activity. To identify structural changes underlying the larger vesicle pool, we used serial blockface electron microscopy of endbulbs from control and noise-exposed mice to quantitatively assess synaptic morphology. We observed no differences in the juxtapositional area between endbulbs and bushy cells, nor in the number or density of active zones and postsynaptic densities. Images of endbulb terminals were significantly darker after noise exposure, indicating an increase in the density of synaptic vesicles. These results suggest that moderate noise exposure induces an activity-dependent increase in presynaptic vesicle numbers, consistent with the observed physiological changes in neurotransmitter release. This work sets the stage for high-resolution studies to quantify docked and reserve vesicles.

**Significance statement:** Noise exposure is a fact of everyday life, and it is important to understand how noise affects function in the auditory pathway in the brain to understand the full consequences of noise exposure. Electrophysiological experiments indicate that noise triggers a homeostatic increase in the releasable pool of vesicles at auditory nerve synapses. We examined the cellular basis for this change using serial blockface electron microscopy of auditory nerve synapses with and without noise exposure. We reconstructed a number of bushy cells and their presynaptic auditory nerve terminals. After noise exposure, there was no significant increase in the area of synaptic contact or the number or density of synaptic release sites. There was an increase in the number of vesicles near release sites, which may account for the physiological changes. These results emphasize the importance of detailed anatomical studies to study the effects of noise exposure and thus determine the best mechanistic approach for therapies and treatments of noise-induced hearing loss.

## Introduction

Prolonged noise exposure (NE) alters auditory function, but the extent of cellular and ultrastructural changes it induces in the central auditory pathway remains unclear (Ayeni et al., 2025). Previous studies have documented the effects of hearing loss, such as caused by deafferentation, cochlear ablation, tympanectomy, or genetic mutation, which can lead to synaptic degeneration and reorganization in the cochlear nucleus (Mendoza Schulz et al. 2014; Clarkson et al. 2016; Rubel et al. 1990; Ryugo et al. 1997). In contrast, the effects of increased sound-driven activity are much less studied, even though moderate, non-traumatic NE is a very common experience in modern life.

To understand this issue, we focus on endbulbs of Held, which are large axosomatic terminals formed by auditory nerve afferents onto bushy cells in the anteroventral cochlear nucleus (AVCN) (Nó 1981). These synapses are specialized for precise timing and high-frequency transmission (Ryugo et al. 1997; Oertel 1999). Endbulbs change physiological properties in response to non-traumatic NE, with decreased probability of vesicle release and increased readily releasable pool (RRP) of vesicles (Ngodup et al. 2015; Wong and Xu-Friedman 2022). These changes appear to be homeostatic, preserving postsynaptic spiking despite high rates of presynaptic activation (Ngodup et al. 2015). Anatomically, prolonged NE leads to larger areas immunopositive for VGluT1 surrounding bushy cells, most likely indicating increased numbers of vesicles in the presynaptic terminals of auditory nerve afferents. It is not clear how this relates to the RRP, which could expand in a number of ways, including overall larger terminals, more release sites, or more releasable vesicles at individual release sites. Distinguishing these alternatives requires better understanding of synaptic ultrastructure.

To address this, we used serial blockface electron microscopy (SBFEM) to visualize the ultrastructure of endbulb terminals in mice exposed to prolonged moderate-level noise. We reconstructed individual synaptic terminals and quantitatively assessed juxtapositional membrane area (JPA), number and density of release sites, and vesicle density. Our findings suggest that moderate NE induces minor changes in size and number of release sites, but significantly increases vesicle density. These results improve our understanding of how auditory nerve terminals adapt to increased acoustic activity.

## Methods

### Animals and NE

All procedures were performed according to approved institutional animal care and use guidelines. CBA/CaJ mice (Jackson Labs strain #000654) were bred in the animal facility in a quiet room. Control mice remained in the quiet room, and NE mice were kept in a separate room beginning at P21 with a Fostex FT28D speaker placed on top of the cage driven by an ACO Pacific white noise generator (model 3025). The noise generator delivers a signal that is flat (within 3 dB) between 1 and 35 kHz, and rolls off at 4 dB/octave above 40 kHz.

Sound levels in the quiet room and in the noise cages were measured using a Larson-Davis sound level meter at 1/3 octave intervals between 1 and 20 kHz. In the quiet room, sound levels averaged 27 dB SPL (range 21 to 38 dB SPL), and in the NE cages, the average level was 81 dB SPL near the speaker, and 74 dB SPL at the far end of the cage (range 64 to 86 dB SPL). Mice were held in noise for 7 d, when physiological assays of vesicle pool size show robust changes (Wong and Xu-Friedman 2022).

### Tissue fixation and sectioning

At P28, 3 control and 3 NE mice were transcardially perfused with 2.5% glutaraldehyde and 4% paraformaldehyde in 0.1 M sodium cacodylate buffer (pH 7.2). Brains were extracted and the cochlear nucleus dissected. Thick sections (150 µm) were cut on a Campden Integraslicer 7550 MM, placed in vials in fix and shipped to Renovo Neural (Cleveland, OH) for staining, embedding, and imaging.

### Staining and embedding

Sections were washed in cacodylate buffer, incubated in 0.1% tannic acid for 30 min, washed in cacodylate buffer, incubated in 2% OsO_4_ plus 1.5% potassium ferrocyanide in cacodylate buffer for 2 hr on ice, washed with water, incubated for 30 min in 1% thiocarbohydrazide (TCH) at 60°C, washed with water, incubated in 2% OsO_4_ for 1 hr, washed with water, incubated overnight in 1% uranyl acetate, washed with water, incubated for 30 min at 60°C in 0.7% lead nitrate in 0.4% aspartic acid (pH 5.5, adjusted with KOH), and washed with water. Sections were dehydrated through an alcohol series (50%, 75%, 85%, 95%, 100%) followed by propylene oxide, and embedded in Epon (47% Epon 812, 21% DDSA, 30% NMA, 2% DMP30).

### SBFEM imaging

Sample quality was assessed by inspecting thin sections, and single blocks from control and NE mice were selected for serial imaging and analysis. The control series was imaged using a Zeiss Sigma VP, consisting of 700 serial images, each with an area of 67.37 µm x 67.37 µm, with a resolution of 6.6 nm/pixel and 70 nm/slice. The NE series was imaged using a FEI Teneo VolumeScope, consisting of four 61.44 µm x 61.44 µm tiles with a resolution of 6 nm/pixel and 65 nm/slice. The tiles in the NE series were stitched together manually using a custom-written program in Wavemetrics Igor.

### Annotation and reconstruction

The control series was annotated using Reconstruct (Fiala 2005), and the NE series was annotated using WebKnossos (Boergens et al. 2017). JPA was quantified as the area where membranes of the endbulb and bushy cell soma were within a distance of 0.15 µm of each other. For control data in Reconstruct, this was quantified by summing segments of bushy cell soma traces within 0.15 µm of a given endbulb and multiplying by section thickness to compute JPA. For NE data in webKnossos, annotated structures were exported as triangulated meshes (.stl files), and JPA was computed as the summed area of triangles on the bushy cell that were within 0.15 µm of any triangle on a given endbulb. Average pixel brightness in the terminal was assessed using the measure tool in ImageJ over an area of 111 x 111 pixels. The brightness was normalized to the average brightness over an identical area in a nearby bushy cell nucleus.

### Statistical Analysis

Synaptic measurements in Fig. 4 were tested for normality using the Shapiro-Wilk test. Summary statistics for normally-distributed data (release site count) are reported as mean ± standard error of the mean (SEM), and populations were compared using Student’s *t* tests. Summary statistics for non-normally-distributed data (JPA, release site density) are reported as median ± median absolute deviation (MAD), and populations were compared using Mann Whitney U tests.

## Results

### Identification of bushy cells and endbulbs

To identify ultrastructural contributions to changes in vesicle pools, we generated SBFEM datasets of the cochlear nucleus in control and NE mice, and annotated bushy cells, endbulbs, and individual release sites. Bushy cells were identified based on soma size and shape (round or ovoid, diameter ∼15 µm), nucleus position and shape (somewhat offset from the center of the soma), cytoplasm (large volume relative to nucleus, numerous ER rosettes and mitochondria), and a single, highly branched dendrite (Fig. 1A, D) (Cant and Morest 1979; Brawer et al. 1974; Lauer et al. 2013; Jing et al. 2024; Spirou et al. 2023). The dendrites of bushy cells were not well-captured in the series, so we restricted annotation of synapses to those made directly on the bushy cell somata.

**Figure 1.**
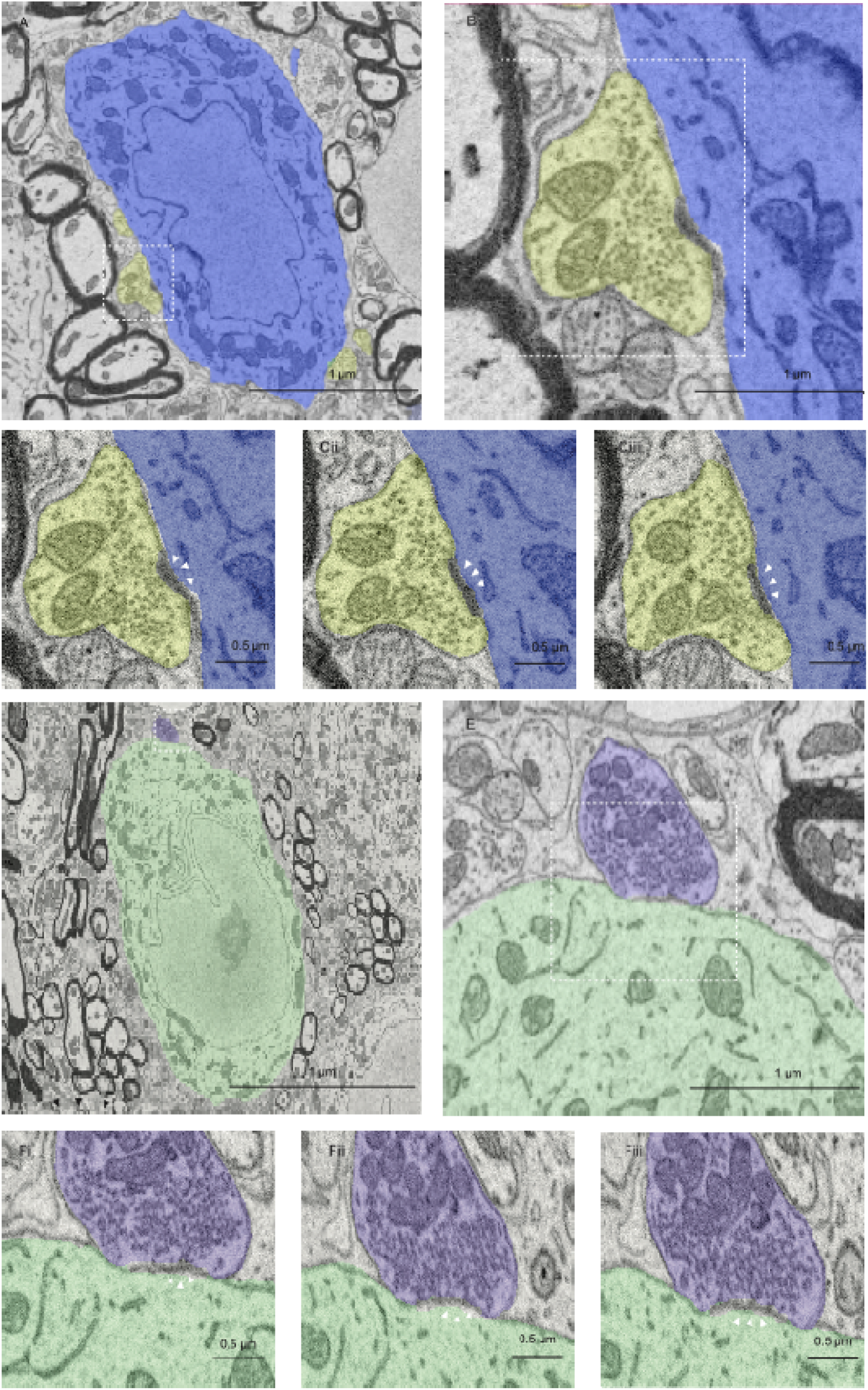
SBFEM of bushy cells and endbulbs in the anteroventral cochlear nucleus of control and NE mice. **A–C**, Bushy cell (blue) and endbulb (yellow) from the control series. The dashed box in **A** is magnified in **B**. The release site in **B** is shown in serial sections in **Ci–iii**. Arrowheads marks the PSD. **D–F**, Bushy cell (green) and endbulb (purple) from the NE series. The dashed box in **D** is magnified in **E**. The release site in **E** is shown in serial sections in **Fi–iii**. Arrowheads marks the PSD.

Endbulbs were identified by curved axosomatic postsynaptic densities (PSDs) flanked by cisternae (Fig. 1B, E). Vesicle shape can be a useful criterion for recognizing endbulbs in EM, because excitatory synapses usually have large, round, clear vesicles. However, the resolution was not adequate to distinguish vesicle shape. As synapses were reconstructed and inspected in 3D, we verified that they had overall complex morphology with multiple axosomatic terminal swellings, electron lucent cytoplasm, and characteristic extended extracellular spaces (Lauer et al. 2013; Limb and Ryugo 2000; O’Neil et al. 2011). Release sites were identified by a PSD across from a presynaptic cluster of vesicles (Fig. 1C, F).

### Quantitative comparisons between control and NE

We reconstructed bushy cells and endbulbs, and annotated PSDs in control and NE samples (Fig. 2). There were 6 bushy cells and 26 endbulbs in the control sample, and 4 bushy cells with 22 endbulbs in the NE sample. To detect changes in overall size or PSD number, it was important to study endbulbs that were completely contained in the series. There were 20 complete endbulbs in the control sample and 9 in the NE sample. We quantified sizes of endbulbs using the JPA, which varied considerably (Fig. 3), consistent with the large variability of synaptic currents between different CBA/CaJ endbulbs recorded in electrophysiology experiments (for example: (Chanda and Xu-Friedman 2010; Yang and Xu-Friedman 2012; Xie and Manis 2017; Wong and Xu-Friedman 2022). The median JPA was larger after NE, but the increase was not significant (control: 22.6 ± 15.5 µm^2^, 20 endbulbs; NE: 48.8 ± 26.7 µm^2^, 9 endbulbs; *p* = 0.10, one-tailed Mann-Whitney U; Fig. 4A). We also considered changes in the JPA size relative to the surface area of BC soma in the series, which allowed us to include values from incomplete endbulbs. The relative JPA was not significantly larger after NE (control: 3.0 ± 2.2%, 26 endbulbs; NE: 2.4 ± 1.5%, 22 endbulbs; *p* = 0.88, one-tailed Mann-Whitney U; Fig. 4B).

**Figure 2.**
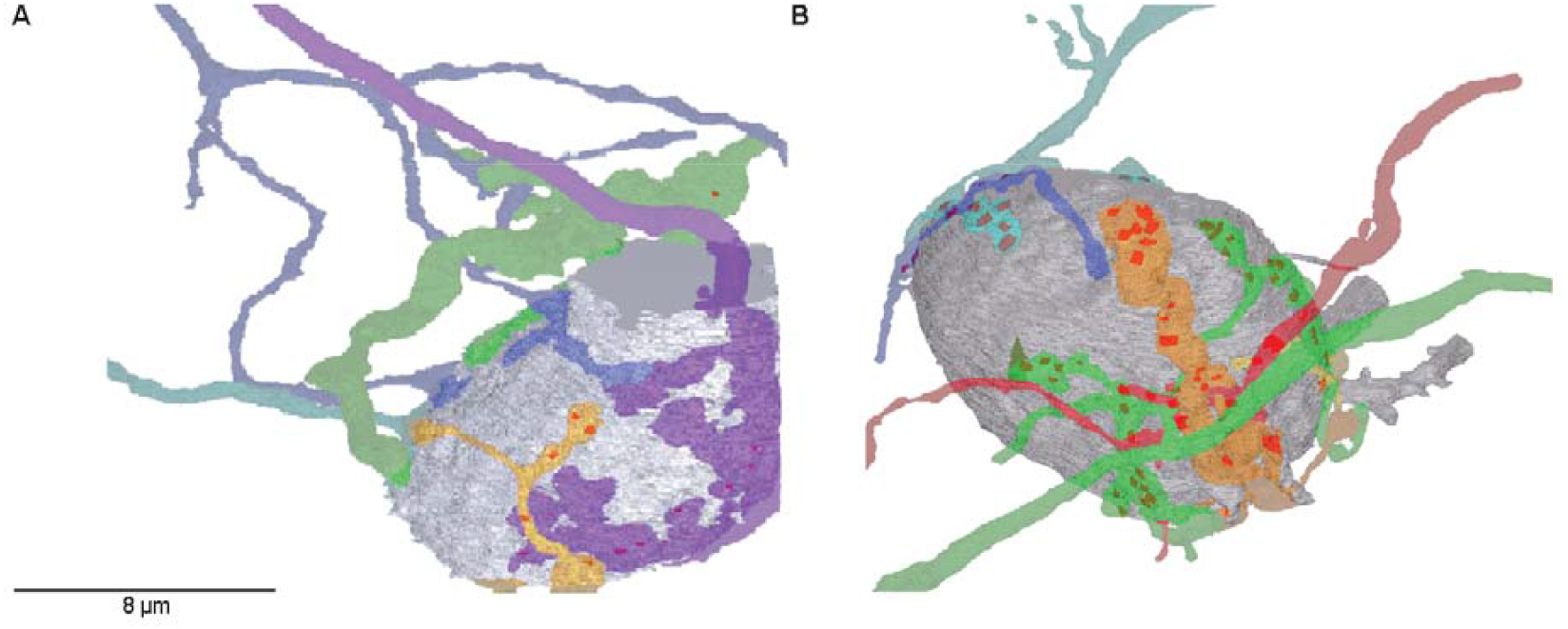
Representative 3D reconstructions of bushy cells and endbulbs from control (left) and NE (right) conditions. Bushy cell somata are shown in gray, and individual auditory nerve fibers are shown in distinct colours. Darker patches on endbulbs mark the locations of PSDs.

**Figure 3.**
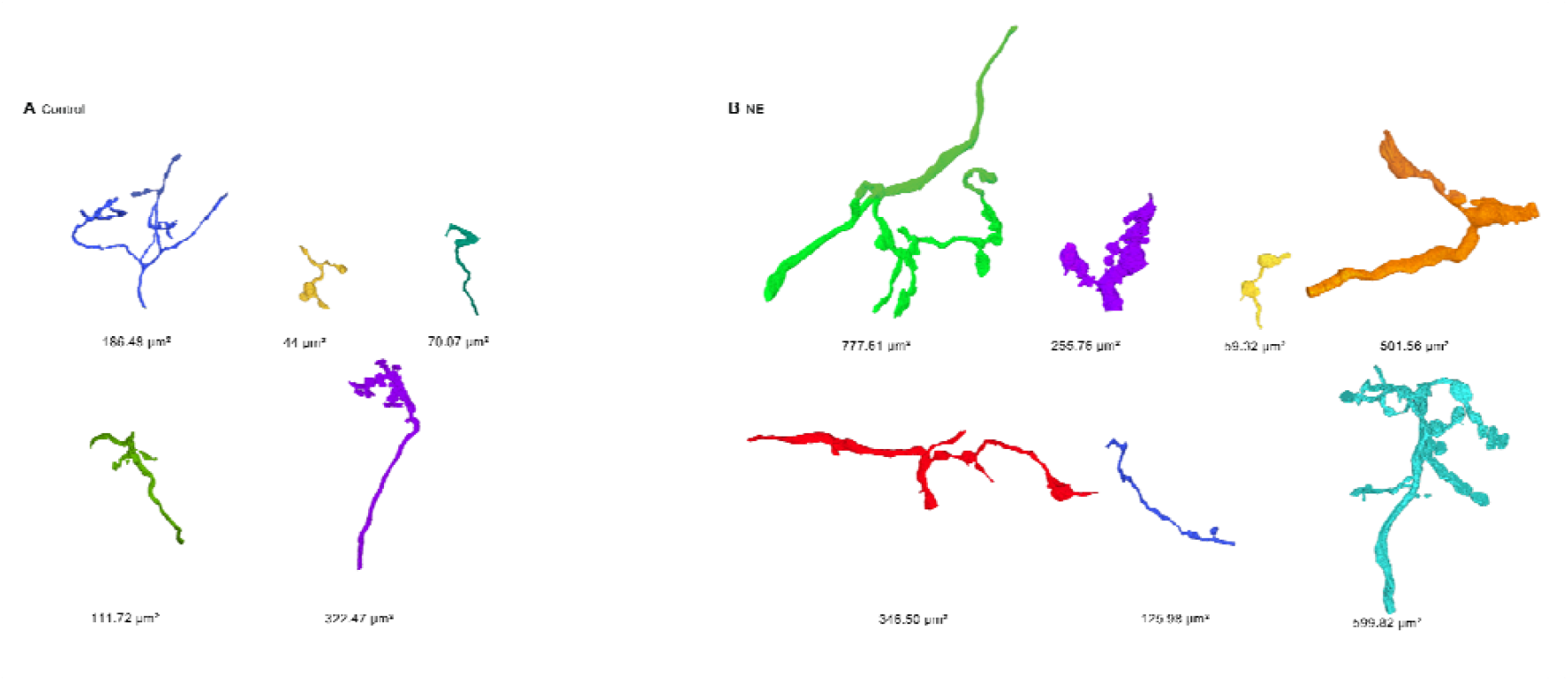
Endbulbs formed onto representative bushy cells from (A) control and (B) NE mice. Each endbulb is rotated to optimize the viewing angle. JPA is quantified below each endbulb. Different colors are used for each endbulb.

**Figure 4.**
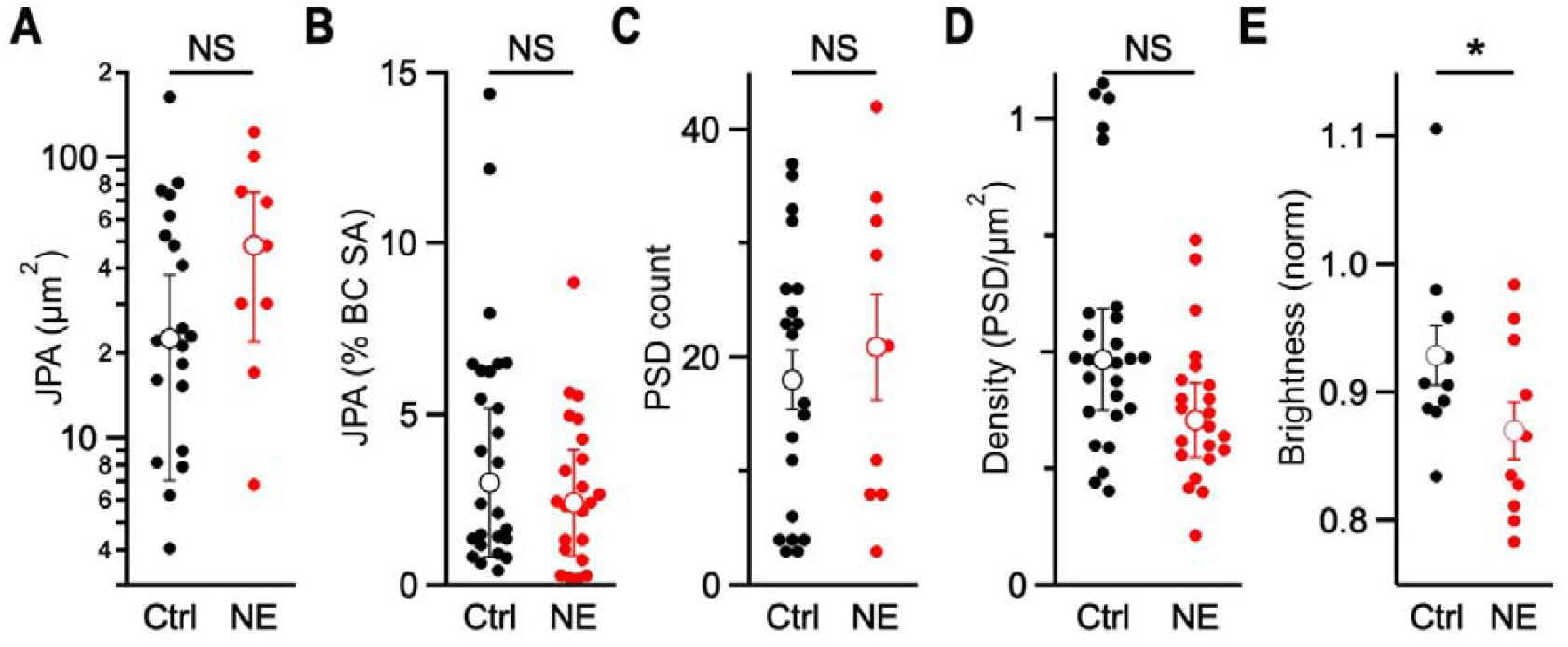
Quantification of synaptic characteristics in reconstructed endbulbs. There were no significant increases in (**A**) JPA (*p* = 0.10), relative JPA (*p* = 0.88), (**C**) PSD count (*p* = 0.35), or (**D**) PSD density (*p* = 0.99) after NE. There was a significant decrease in (**E**) normalized pixel brightness (*p* = 0.046) in the presynaptic terminal after NE, which is consistent with an increase in the number of vesicles.

The RRP could also enlarge by increasing the number of release sites, with or without a change in JPA. We directly examined release sites by quantifying the number of PSDs in control and NE endbulbs. In the complete endbulbs, there was no significant change in the absolute number of PSDs after NE (control: 18.1 ± 2.6, 20 endbulbs; NE: 20.9 ± 4.7, 9 endbulbs; *p* = 0.29, one-tailed *t* test; Fig 4C). Furthermore, we also evaluated changes in PSD density (#PSDs/µm^2^). This parameter is relative, so we could include values from all endbulbs irrespective of whether they were completely contained in the series. PSD density also showed no significant increase after NE (control: 0.48 ± 0.11 PSDs/µm^2^, 26 endbulbs; NE: 0.36 ± 0.08 PSDs/µm^2^, 22 endbulbs; *p* = 0.99, one-tailed Mann-Whitney U; Fig. 4D). Thus, neither overall growth of endbulbs nor insertion of new PSDs accounts for the increased RRP seen after NE.

Finally, the RRP could increase after NE by increasing the number of vesicles at each release site. The resolution of SBFEM was not adequate for distinguishing docked and undocked vesicles or for counting vesicles. Instead, we compared control and NE endbulbs by measuring average pixel brightness within presynaptic terminals. To correct for exposure differences between images, we normalized values to average brightness of the nucleus of the postsynaptic cell. We found that brightness was significantly lower in presynaptic terminals after NE (control 0.93 ± 0.02, 10 endbulbs; NE 0.87 ± 0.02, 10 endbulbs; *p* = 0.046, one-tailed *t* test; Fig. 4E).

Vesicles appear black in these images, so the decrease in brightness is consistent with an increased number of vesicles after NE. It will be of great interest to quantify changes in numbers of docked and reserve vesicles in material of higher resolution in the future.

## Discussion

This study used SBFEM to examine whether moderate, non-traumatic NE produces measurable ultrastructural changes in auditory nerve endbulbs contacting bushy cells. Previous electrophysiological work has shown that similar exposure conditions lead to decreased release probability and an expanded readily releasable pool (RRP) of vesicles (Ngodup et al. 2015; Wong and Xu-Friedman 2022). We proposed three potential structural mechanisms that could underlie these changes: overall endbulb growth with increased numbers of release sites, greater density of release sites within the same contact area, or an increase in vesicles associated with existing active zones. No significant differences were detected in JPA, PSD count, or PSD density between control and NE samples. The only consistent change was a decrease in normalized pixel brightness within NE terminals, consistent with an increase in vesicle density.

Earlier single-section EM work reported increases in presynaptic perimeter and area after NE (Ngodup et al. 2015), suggesting that terminals enlarge. The present reconstructions refine that view, suggesting that the contact area with the bushy cell soma does not expand significantly. That would imply that the prior increases in cross-sectional perimeter and area could reflect thickening of terminals more than growth in the area of contact between endbulb and bushy cell, perhaps to accommodate more mitochondria and synaptic vesicles. This raises the question of how other structures in the AVCN change to make way for larger endbulbs, which will require analysis of additional structures in these series.

To assess changes in vesicle numbers in endbulbs, we compared the brightness of images of presynaptic terminals, which showed a significant decrease in NE endbulbs, consistent with increased numbers of vesicles. It would be highly valuable to confirm this result with direct counts. The resolution in SBFEM images was too low to quantify docked vesicles (which presumably reflect the RRP) or vesicles further from the active zone (which presumably reflect reserve vesicles) (Chicurel and Harris 1992; Schikorski and Stevens 1997; Xu-Friedman et al. 2001; Kaeser and Regehr 2017). Furthermore, tissue sections are destroyed during SBFEM imaging, so it is not possible to return to a section once the images are disrupted. This makes it difficult to capture complete endbulbs.

Another approach may be to use ATUMtome, which provides high stability for thin sections collected on tape, which are then imaged using SEM (Schalek et al. 2012). This approach would allow first gathering overviews of entire endbulb–bushy cell contacts at low resolution, followed by imaging individual release sites at high resolution to assess vesicle shape or count individual vesicles that are docked near release sites or within reserve pools. In addition, ATUMtome could reduce a problem of SBFEM and serial electron microscopy more generally that it is very difficult to know that an area initially selected for imaging will include structures of interest as imaging proceeds. In this study, the control series fortuitously contained several bushy cell somata and endbulbs in their entirety, while the NE series had fewer. ATUMtome would make it easier to optimize what to image and to adjust the area to be imaged to follow important structures. This may improve sample sizes making it feasible to investigate additional issues, such as how quickly synapses change in noise and recover after return to normal acoustic conditions.

Previous work also assayed changes in endbulb anatomy using immunohistochemistry against VGluT1, which labels glutamatergic vesicles in puncta surrounding bushy cell somata (Lauer et al. 2013). After NE, immunolabelled puncta increased in size, but not in number (Ngodup et al. 2015; Wong and Xu-Friedman 2022). The pixel brightness metric used here (Fig. 4E) indicates that vesicle density increased after NE, which would appear brighter in fluorescence images of VGluT1 immunolabelling. Brighter labelling would likely increase the volume of presynaptic terminal in which fluorescence is detectable, which would translate to larger puncta. Thus, the changes in ultrastructure seen here are consistent with previous measurements using immunohistochemistry. Furthermore, EM provides much greater insight into structural modifications than immunohistochemistry.

It will be important to expand this study to consider the consequences of conductive hearing loss (CHL), which decreases auditory nerve activity. CHL is extremely common in humans, and may contribute to central auditory processing disorders through cellular changes that aren’t well understood (Whitton and Polley 2011). If the central effects of CHL are opposite to NE quantified in this study, one prediction could be that the number and density of PSDs would remain unchanged, while the number of vesicles per PSD decreases. After a week of CHL, the RRP assessed using electrophysiology decreases at endbulbs, and there is a parallel decrease in the size of VGluT1-immunopositive puncta (Zhuang et al. 2017; Wong and Xu-Friedman 2022). Other studies have examined the consequences of various forms of deafness. Endbulbs of mice with knockouts of Bassoon have smaller RRPs and similar numbers of active zones (Mendoza Schulz et al. 2014), suggesting fewer vesicles per PSD. Mice with progressive early-onset hearing loss have smaller endbulbs, and largely similar appositional area (Connelly et al. 2017; Ayeni et al. 2025). PSD size increases in some models of CHL and deafness (Clarkson et al. 2016; Ryugo et al. 1997; Lee et al. 2003), which might be expected to correlate with increased RRP size. Thus, there is room for closer comparison of endbulbs from normal, NE, and CHL mice to understand how changes in activity influence synaptic properties. In this way, our work provides an important step for identifying the anatomical basis through which acoustic experience shapes auditory brainstem plasticity under conditions of increased, rather than decreased, activity.

## Acknowledgements

The authors thank Connor Cook for assistance with histology, Austin D’Angelo, Treefa Shwani, Jarrod Lynch, Daysia Augustin, and Aaron Lee for advice about annotation. This work was supported by National Institutes of Health grants R01 DC015508 (MXF) and the George T. Nager endowment (AML).

## References

Ayeni, Femi E., Michael A. Muniak, and David K. Ryugo. 2025. “Effect of Sound Amplification on Central Auditory Plasticity: Endbulb of Held as a Substrate.” Brain Sciences 15 (8): 888. 10.3390/brainsci15080888.

Boergens, Kevin M., Manuel Berning, Tom Bocklisch, et al. 2017. “webKnossos: Efficient Online 3D Data Annotation for Connectomics.” Nature Methods 14 (7): 7. 10.1038/nmeth.4331.

Brawer, J. R., D. K. Morest, and E. C. Kane. 1974. “The Neuronal Architecture of the Cochlear Nucleus of the Cat.” The Journal of Comparative Neurology 155 (3): 251–300. 10.1002/cne.901550302.

Cant, N. B., and D. K. Morest. 1979. “The Bushy Cells in the Anteroventral Cochlear Nucleus of the Cat. A Study with the Electron Microscope.” Neuroscience 4 (12): 1925–45. 10.1016/0306-4522(79)90066-6.

Chanda, Soham, and Matthew A. Xu-Friedman. 2010. “A Low-Affinity Antagonist Reveals Saturation and Desensitization in Mature Synapses in the Auditory Brain Stem.” Journal of Neurophysiology 103 (4): 1915–26. 10.1152/jn.00751.2009.

Chicurel, M. E., and K. M. Harris. 1992. “Three-Dimensional Analysis of the Structure and Composition of CA3 Branched Dendritic Spines and Their Synaptic Relationships with Mossy Fiber Boutons in the Rat Hippocampus.” The Journal of Comparative Neurology 325 (2): 169–82. 10.1002/cne.903250204.

Clarkson, Cheryl, Flora M. Antunes, and Maria E. Rubio. 2016. “Conductive Hearing Loss Has Long-Lasting Structural and Molecular Effects on Presynaptic and Postsynaptic Structures of Auditory Nerve Synapses in the Cochlear Nucleus.” The Journal of Neuroscience: The Official Journal of the Society for Neuroscience 36 (39): 10214–27. 10.1523/JNEUROSCI.0226-16.2016.

Connelly, Catherine J., David K. Ryugo, and Michael A. Muniak. 2017. “The Effect of Progressive Hearing Loss on the Morphology of Endbulbs of Held and Bushy Cells.” Hearing Research 343 (January): 14–33. 10.1016/j.heares.2016.07.004.

Fiala, J. C. 2005. “Reconstruct: A Free Editor for Serial Section Microscopy.” Journal of Microscopy 218 (1): 52–61. 10.1111/j.1365-2818.2005.01466.x.

Jing, Junzhan, Ming Hu, Tenzin Ngodup, et al. 2024. “Molecular Logic for Cellular Specializations That Initiate the Auditory Parallel Processing Pathways.” bioRxiv: The Preprint Server for Biology, October 6, 2023.05.15.539065. 10.1101/2023.05.15.539065.

Kaeser, Pascal S., and Wade G. Regehr. 2017. “The Readily Releasable Pool of Synaptic Vesicles.” Current Opinion in Neurobiology 43 (April): 63–70. 10.1016/j.conb.2016.12.012.

Lauer, Amanda M., Catherine J. Connelly, Heather Graham, and David K. Ryugo. 2013. “Morphological Characterization of Bushy Cells and Their Inputs in the Laboratory Mouse (Mus Musculus) Anteroventral Cochlear Nucleus.” PLOS ONE 8 (8): e73308. 10.1371/journal.pone.0073308.

Lee, Daniel J., Hugh B. Cahill, and David K. Ryugo. 2003. “Effects of Congenital Deafness in the Cochlear Nuclei of Shaker-2 Mice: An Ultrastructural Analysis of Synapse Morphology in the Endbulbs of Held.” Journal of Neurocytology 32 (3): 229–43. 10.1023/B:NEUR.0000010082.99874.14.

Limb, Charles J., and David K. Ryugo. 2000. “Development of Primary Axosomatic Endings in the Anteroventral Cochlear Nucleus of Mice.” JARO: Journal of the Association for Research in Otolaryngology 1 (2): 103–19. 10.1007/s101620010032.

Mendoza Schulz Alejandro,, Zhizi Jing, Juan María Sánchez Caro, et al. 2014. “Bassoon-Disruption Slows Vesicle Replenishment and Induces Homeostatic Plasticity at a CNS Synapse.” The EMBO Journal 33 (5): 512–27. 10.1002/embj.201385887.

Ngodup, Tenzin, Jack A. Goetz, Brian C. McGuire, Wei Sun, Amanda M. Lauer, and Matthew A. Xu-Friedman. 2015. “Activity-Dependent, Homeostatic Regulation of Neurotransmitter Release from Auditory Nerve Fibers.” Proceedings of the National Academy of Sciences 112 (20): 6479–84. 10.1073/pnas.1420885112.

Nó Rafael Lorente de. 1981. The Primary Acoustic Nuclei. Raven Press.

Oertel, D. 1999. “The Role of Timing in the Brain Stem Auditory Nuclei of Vertebrates.” Annual Review of Physiology 61: 497–519. 10.1146/annurev.physiol.61.1.497.

O’Neil Jahn N., Catherine J. Connelly, Charles J. Limb, and David K. Ryugo. 2011. “Synaptic Morphology and the Influence of Auditory Experience.” Hearing Research 279 (1–2): 118–30. 10.1016/j.heares.2011.01.019.

Rubel, E. W., R. L. Hyson, and D. Durham. 1990. “Afferent Regulation of Neurons in the Brain Stem Auditory System.” Journal of Neurobiology 21 (1): 169–96. 10.1002/neu.480210112.

Ryugo, D. K., T. Pongstaporn, D. M. Huchton, and J. K. Niparko. 1997. “Ultrastructural Analysis of Primary Endings in Deaf White Cats: Morphologic Alterations in Endbulbs of Held.” The Journal of Comparative Neurology 385 (2): 230–44. 10.1002/(sici)1096-9861(19970825)385:2%253C230::aid-cne4%253E3.0.co;2-2.

Schalek, R, A Wilson, J Lichtman, et al. 2012. “ATUM-Based SEM for High-Speed Large-Volume Biological Reconstructions.” Microscopy and Microanalysis 18 (S2): 572–73. 10.1017/S1431927612004710.

Schikorski, T., and C. F. Stevens. 1997. “Quantitative Ultrastructural Analysis of Hippocampal Excitatory Synapses.” The Journal of Neuroscience: The Official Journal of the Society for Neuroscience 17 (15): 5858–67. 10.1523/JNEUROSCI.17-15-05858.1997.

Spirou, George A., Matthew Kersting, Sean Carr, et al. 2023. “High-Resolution Volumetric Imaging Constrains Compartmental Models to Explore Synaptic Integration and Temporal Processing by Cochlear Nucleus Globular Bushy Cells.” eLife 12 (June): e83393. 10.7554/eLife.83393.

Whitton, Jonathon P., and Daniel B. Polley. 2011. “Evaluating the Perceptual and Pathophysiological Consequences of Auditory Deprivation in Early Postnatal Life: A Comparison of Basic and Clinical Studies.” Journal of the Association for Research in Otolaryngology 12 (5): 535–47. 10.1007/s10162-011-0271-6.

Wong, Nicole F., and Matthew A. Xu-Friedman. 2022. “Time Course of Activity-Dependent Changes in Auditory Nerve Synapses Reveals Multiple Underlying Cellular Mechanisms.” The Journal of Neuroscience 42 (12): 2492–502. 10.1523/JNEUROSCI.1583-21.2022.

Xie, Ruili, and Paul B. Manis. 2017. “Synaptic Transmission at the Endbulb of Held Deteriorates during Age□ related Hearing Loss.” The Journal of Physiology 595 (3): 919–34. 10.1113/JP272683.

Xu-Friedman, M. A., K. M. Harris, and W. G. Regehr. 2001. “Three-Dimensional Comparison of Ultrastructural Characteristics at Depressing and Facilitating Synapses onto Cerebellar Purkinje Cells.” The Journal of Neuroscience: The Official Journal of the Society for Neuroscience 21 (17): 6666–72. 10.1523/JNEUROSCI.21-17-06666.2001.

Yang, Hua, and Matthew A. Xu-Friedman. 2012. “Emergence of Coordinated Plasticity in the Cochlear Nucleus and Cerebellum.” The Journal of Neuroscience: The Official Journal of the Society for Neuroscience 32 (23): 7862–68. 10.1523/JNEUROSCI.0167-12.2012.

Zhuang, Xiaowen, Wei Sun, and Matthew A. Xu-Friedman. 2017. “Changes in Properties of Auditory Nerve Synapses Following Conductive Hearing Loss.” The Journal of Neuroscience 37 (2): 323–32. 10.1523/JNEUROSCI.0523-16.2016.

